# Closed-loop error damping in human BCI using pre-error motor cortex activity

**DOI:** 10.64898/2026.02.25.707999

**Authors:** Camille Gontier, William Hockeimer, Nicolas G. Kunigk, Edgar Canario, Linnea J. Endsley, John E. Downey, Jeffrey M. Weiss, Brian Dekleva, Jennifer L. Collinger

## Abstract

Intracortical brain-computer interfaces (BCIs) are used to decode motor intent from neural population activity; their main clinical application is to restore function for individuals with motor or communication deficits. However, when trying to reconstruct movement trajectories, such as in computer cursor control, even state-of-the-art decoders fall short of able-bodied performance during online BCI control. This calls for alternative approaches to improve the usability of motor BCIs. Here, we leveraged an error signal, i.e. a neural correlate of faulty motor control that can be detected from neural activity. By detecting this error signal in parallel to performing movement decoding, it is possible to perform error modulation, i.e. real-time error detection and correction during a closed-loop motor BCI task. We analyzed data from four individuals with upper limb impairment due to cervical spinal cord injury who each used an intracortical BCI to perform a continuous cursor control task with visual feedback. A classifier was trained to detect the error signal and was used to perform online error detection during BCI control to limit ongoing errors (defined as movement of the controller away from its target) without requiring any specific action from the participants. Our contribution is three-fold. First, we show that the error signal has a pre-error component. Cortical activity was already significantly modulated before the onset of the kinematically-defined error, theoretically allowing for earlier detection. Second, we show that error modulation significantly improves performance during online BCI control of cursor kinematics. Finally, we show that the error signal can be robustly leveraged across contexts, as error modulation improves performance in more complex motor tasks (involving for instance grasp and drag actions) or other environments without task-specific calibration. Overall, our results suggest that the error signal can be robustly disentangled from motor intent in cortical activity, and that even a simple linear classifier can enable error modulation in parallel to a continuous kinematic decoder, yielding more reliable and accurate BCI control.

## 1 Introduction

In recent years, brain-computer interfaces (BCIs) have shown promise in restoring motor or communication ability that has been lost or impaired due to injury or disease [1–4]. BCIs use neural recordings, often from motor cortex, to control a computer cursor or assistive devices, thus circumventing the damaged or missing connection between the brain and the limb. State-of-the-art methods have allowed BCI study participants to efficiently access a computer [5–8] and even write letters through imagined handwriting [9].

However, performance in these studies often falls short of able-bodied performance as measured, for instance, by the rate of target selection or by the shape of the decoded trajectories, which can appear wobbly and off-target. Moreover, the performance and precision of BCI control can be highly variable, even within a single session [7, 9]. Different phenomena have been proposed to account for the imperfect and variable decoding performances, including having a limited number of recoding channels [10], recording instabilities [11, 12], the decoder model not fully capturing neural variance [13], or changes in the tuning properties of recorded units [14, 15]. Recent results have highlighted the potential of non-linear techniques (including neural-network-based approaches) to improve the performance of BCI [13]. Although these non-linear methods show promising improvements in decoding accuracy in offline datasets, they still fall short of able-bodied performance during actual closed-loop BCI control [5]. This is especially problematic as BCI users with motor impairments have been reported to prioritize device accuracy over other characteristics [16].

One possibility to improve BCI performance and usability is the augmentation of existing decoding approaches by performing closed-loop, real-time error detection and modulation [17, 18]. To do so, periods of erroneous control must be detected. This could include epochs in which the controlled cursor is moving away from the target or periods in which a prosthetic hand is moving away from an object that the user wants to grasp. For this detection to be performed during unsupervised BCI control, it needs to be agnostic to the user’s intent and to the position of the target (which are typically unknown to the BCI system). One possible approach is to perform error detection using only the neural data itself. The neural features corresponding to erroneous control can be identified via supervised means, i.e., during a training period where the target and the user’s intents are known. Then, when these neural features are detected in a target-agnostic fashion online, the standard decoding algorithm output should be augmented to perform active error damping or correction. Prior work suggests that error-related activity can be observed in neural recordings. For example, BCIs that rely on scalp electroencephalography (EEG) recordings often make use of error-related potentials, i.e. the neural activity that is time-locked to an error or to a failed trial [19–27]. Using intracranial electrodes, previous work has observed error-related signals in different brain areas, with electrocorticography (ECoG) [28, 29], intracortical microelectrode arrays [18, 30], or intracranial EEG with human participants [31].

Detecting this error signal opens the possibility of performing online error detection and, potentially, correction. Previous studies have shown that error-related information can be detected during BCI control: if an anticipated or ongoing error is detected, the decoding scheme can be updated to prevent the error from increasing or correct it, without any action from the participant [17, 18].

One study in non-human primates (NHPs) [17] performed error detection at a trial level, i.e., detecting if the current trial was going to be successful or not. The macaque was instructed, at each trial, to control a BCI to reach for a specific target on a grid. If neural correlates of an incorrect target selection were identified, the trial time was extended to allow for correction before the end of the trial, yielding a significant performance improvement in terms of bit rate.

Error detection has also been performed continuously in a subsequent study with NHPs [18]. Using data from a 2D finger-positioning BCI task, a classifier was built to identify error periods from sliding windows of neural data where error was defined as an increase in the distance between the target and the finger position. During online BCI control, when an error was detected, the decoded velocity was set to zero to prevent the finger from moving further away from its target. This yielded a significant improvement of the stopping ability of the controller for one of the subjects. Error modulation has also shown promise in a brain-to-text speech neuroprosthesis [32] and for a binary classification task [31]. Here, we aim to extend this prior work to implement error modulation while human BCI users perform more complex, continuous motor control tasks. In our implementation of error modulation, we propose damping the BCI control signal rather than setting it to zero so that the user still receives visual feedback of their actions and can make appropriate corrections.

It is usually assumed that this neural error signal is being driven by the sensory (i.e., visual in the case of a cursor control task) feedback of an ongoing error and that it can only be detected after the onset of an error. Indeed, the neural correlates of erroneous control have either been studied in a time window following the onset of the error [18, 19, 25, 29, 31, 32] or by artificially inducing errors, e.g., by modifying the position of the target or cursor [22, 25, 26, 28]. For online error modulation, it is desirable to detect errors as soon as possible, ideally before the error even starts. Here, we investigated whether neural signatures of errors can be detected before their actual onset is reflected in the decoded kinematics.

Our contribution is three-fold. First, we leveraged data from four human participants performing a cursor control task using an intracortical BCI (section 2.1). In line with previous studies in NHPs, we identified significant differences in the neural population activity of motor cortex between periods of correct and erroneous control and show that a classifier can be trained to detect these errors. Additionally, we extended prior work to show that these changes in neural activity can already be observed in a time window (at least up to 80 ms) before the kinematic error begins, allowing for earlier implementation of error modulation without a decline in classification performance (section 2.2). We quantified the differences in cortical activity between periods of correct and erroneous control and showed that neural subspaces during both types of control are significantly misaligned and that neural activity during erroneous control is characterized by a reduction of its dimensionality (section 2.3). Second, we showed that online error modulation, in which velocity was reduced to 30% of the decoded value when an error was detected, led to a significant improvement of BCI cursor control performance (section 2.4) as evidenced by different quantitative performance improvements for each participant. Finally, we demonstrated that neurally-driven error modulation can generalize across contexts and can be used in more complex BCI applications including a gamified click-and-drag BCI task, or tasks from the recent Cybathlon competition [33] (section 2.5). Taken together, this work identified early neural signatures of BCI errors and improved control by modulating the decoded output. This addresses a significant user priority of delivering more reliable control without compromising performance [16, 34].

## 2 Results

### 2.1 Experimental setting

Data were recorded from four human participants with cervical spinal cord injuries who had intra-cortical microelectrode arrays (Blackrock Microsystems, Inc., Salt Lake City, UT) implanted in their motor cortex as part of an ongoing BCI clinical trial conducted under an FDA Investigational Device Exemption NCT01894802 (Fig. 1A). For participants P2, P3, and P4, we used historical data from a 2D+click BCI control task described in [6]. At the beginning of each trial, a cursor appeared in the middle of the screen and participants were instructed to reach to a target randomly located at one of eight possible positions around the center (*reach* phase), click and hold the target, and move the cursor back to the center (*center* phase). To perform BCI control, a decoder was calibrated each day to predict 2D cursor velocity as well as click and unclick (see Movement decoding for details). Participant C2 performed a slightly different version of the task, i.e. a simpler 2D BCI control task without click or grasp. For all participants, data were analyzed from the reach phase of the BCI control task.

**Figure 1:**
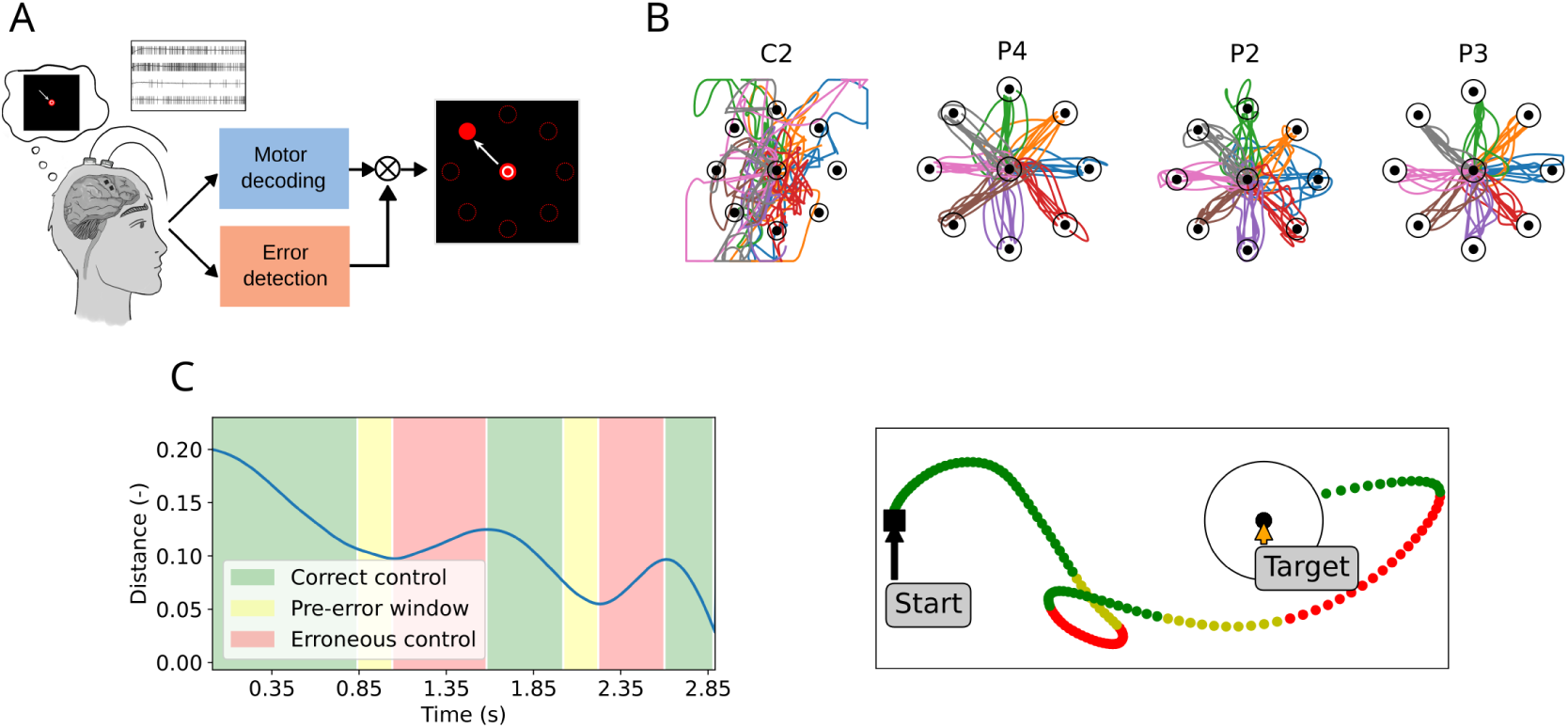
Experimental setting. A: Schematic of the BCI setting. Participants used motor imagery to control the velocity of a cursor on a screen. Right: Schematic of the 2D cursor control center-out task. Each trial starts with the cursor (white circle) at the center of the screen and a random target (red), which participants are instructed to reach. Redrawn from [6]. B: Representative example sessions of cursor control without error modulation for each participant. C: Left: Absolute distance (arbitrary screen units) between target and current cursor position (blue line) for an example trial. Epochs of increasing and decreasing error are marked in red and green, respectively. The pre-error window (in yellow) corresponds to the period of good control immediately preceding the onset of an error. Right: Cursor trajectory for the same trial, represented in the 2D space. The black circle represents the target success radius.

Our decoding approach typically enabled successful acquisition of all targets even without error modulation (Fig. 1B). 3 participants (P2, P3, and P4) had average success rates above 90% (P2: 92.5% *±* 8.8 over 31 sets; P3: 98.2% *±* 4.0 over 18 sets; P4: 98.6% *±* 3.0 over 33 sets), while one participant had lower success rates using the same decoding approach (C2: 36.6% *±* 15.5 over 22 sets), due to their lower number of active electrodes (Supp. Fig. 1). Performance variability across participants and even across trials (as can be seen in the sample trajectories in Fig. 1B), highlights the need for performance improvement, which we aim to accomplish via error modulation.

In line with previous studies [18], we defined periods of correct and erroneous BCI control using the instantaneous distance between the current position of the cursor and the target. Periods of erroneous control were defined as the periods during which this distance between cursor and target is increasing (red in Fig. 1C) instead of decreasing (see section 4.4: Error definition). We then implemented a classifier that computed the probability of an error from online neural recordings.

Before deploying the classifier online, we sought to improve the responsiveness of the classifier which as defined above can only detect an error after the cursor is already deviating away from the ideal path. We explored whether we could leverage the *pre-error* period (yellow in Fig. 1C) and label it as *erroneous control* in the training set, with the goal of detecting errors immediately before their onset. We compare the performance of an error classifier that included this pre-error window to a classifier that only included data after the kinematic error had begun in the next section.

### 2.2 Application of error detection to offline data

Before implementing error modulation during closed-loop real time BCI control, we verified the possibility of detecting erroneous control from neural recordings by training and testing a classifier on offline data. To perform meaningful quantification of neural activity during periods of erroneous control, and to ensure that the dataset used for training the error classifier was sufficiently unbiased, we filtered previous experimental sets to only retain those in which at least 30% of epochs were labeled as *erroneous control* (this threshold was lowered to 25% for participant P4, to account for their better control performance, see Supplementary Table 1 for a detailed list of the used recordings).

A naive Bayes classifier was trained and tested (see section 4.5: Error classification for details) for each session via 8-fold cross-validation (Fig. 2A). Since the datasets are unbalanced (there are usually more periods of correct than of erroneous control), we compared the classifier’s accuracy to that of a greedy classifier which would assign the label corresponding to the most populated class to all samples. For all participants, classification accuracy was higher than this conservative measure of chance-level accuracy (Fig. 2B. *Chance level* vs. *Error only*: one-sided paired t-test. C2: *p <* 0.001; P2: *p <* 0.001; P3: *p* = 0.029; P4: *p* = 0.004; *Chance level* vs. *Error + Pre-error*: one-sided paired t-test. C2: *p <* 0.001; P2: *p <* 0.001; P3: *p* = 0.034; P4: *p* = 0.002.).

**Figure 2:**
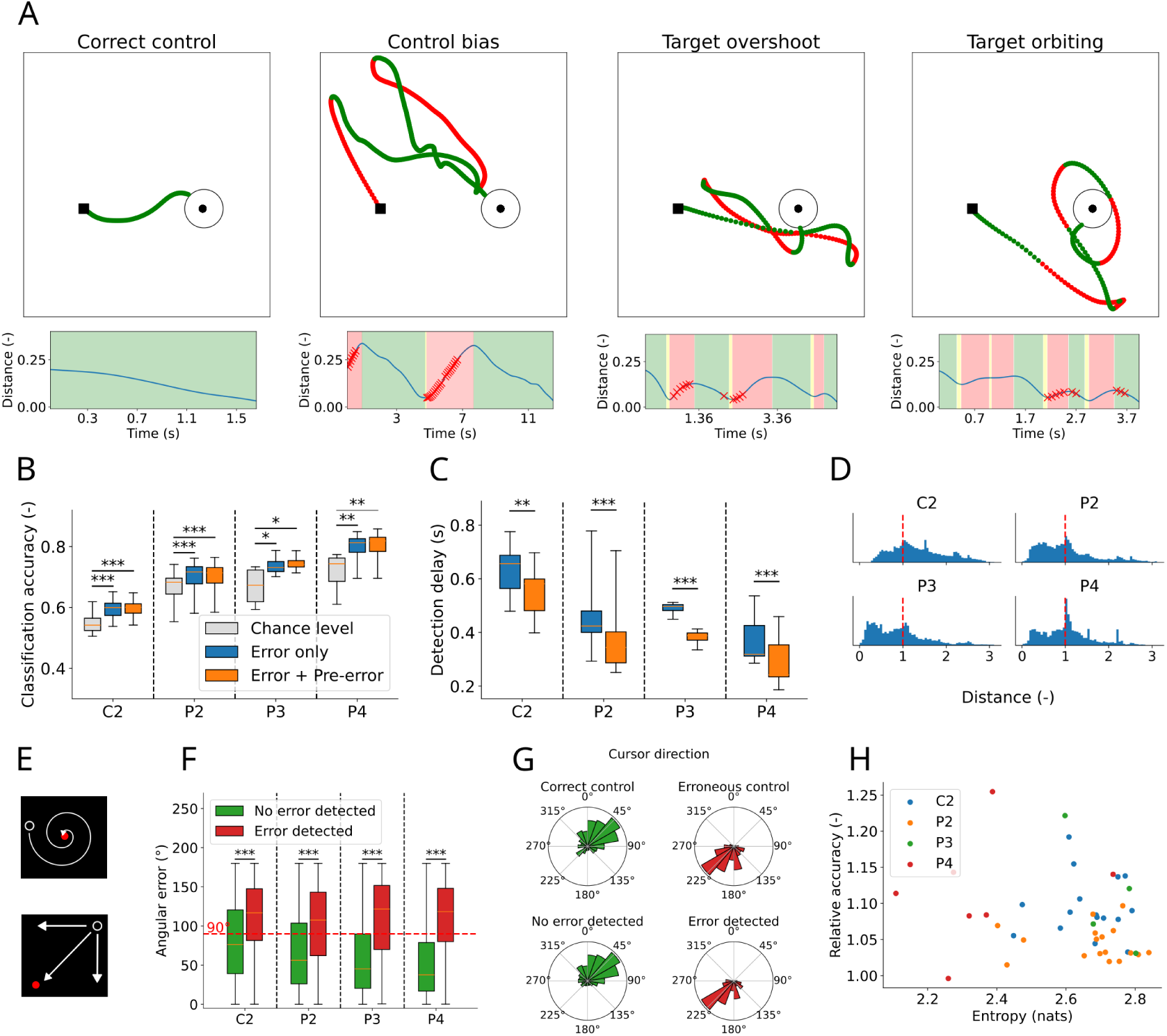
Application of error detection to offline data. A: Top row: Trial examples illustrating different control and failure modes, from left to right: correct control (participant P4), control bias (P4), target overshoot (P2), and target orbiting (P2). Bottom row: Output of the error classifier. Epochs of erroneous control (defined as epochs of increasing distance between cursor and target) are displayed in a red background. Red crosses indicate epochs (held-out during training) at which the classifier detects an error. B: Chance levels (light grey) vs. classification accuracies, without (blue) and with (orange) leveraging the pre-error window for training the classifier across all sessions. C: Mean delay between the onset of erroneous control and its detection by the classifier across all sessions and as a function of whether the pre-error window is leveraged for training the classifier. D: Distribution of the distance between cursor and target during epochs labeled as erroneous control for all sessions. Values are normalized by the distance from the center to the target at the beginning of the trial (dashed red line). E: Examples of non-optimal trajectories that would nonetheless be classified as correct control. Top: the cursor is spiraling towards the target and the distance between them is continuously decreasing. Bottom: whatever the trajectory followed by the cursor, the distance between it and the target decreases (although only the diagonal trajectory is optimal). F: Angular errors during epochs where the classifier does not report an error (green) vs. angular error during epochs where an error is reported (red). G: Histograms of the cursor velocity direction during epochs of correct control (top left), erroneous control (top right), during held-out epochs where the classifier does not (bottom left) or does detect an error (bottom right), for one example session (consisting of several trials) from participant P4. H: Entropy of the cursor velocity direction vs. relative classification accuracy (ratio of accuracy to chance level in Fig. 2B) for all sets.

Moreover, we verified that including the pre-error window in the training of the classifier allowed for a significant reduction in the delay between the onset of faulty control as defined by kinematics and its detection by the classifier (Fig. 2C; C2: *p* = 0.001; P2: *p* =*<* 0.001; P3: *p* = 0.001; P4: *p <* 0.001, one-sided paired t-test) while marginally modifying classification accuracy (no significant differences between “Error only” and “Error + Pre-error” in Fig. 2B. Two-sided paired t-test. C2: *p* = 0.153; P2: *p* = 0.42; P3: *p* = 0.198; P4: *p* = 0.088).

For each time point that was labeled as erroneous, we computed the distance from the target based on the current cursor position. The goal of this analysis was to determine if there was any pattern of error types (e.g. moving incorrectly from the start vs. orbiting near the target) within or across participants. We saw no clear typology of errors during BCI control, but we did observe two peaks to varying degrees in all participants (Fig. 2D): the first peak (distance between 0 and 0.5) corresponding to small corrective movements close to the target, the second one (centered around 1 with a longer tail) to errors in which the cursor was farther away from the target than when the trial started.

As in previous studies (e.g. [18]), the metric that we used to define correct vs. erroneous epochs was based on the distance between the current position of the cursor and the target (and more specifically whether this distance is increasing or decreasing). Although objective (in the sense that it does not require setting an arbitrary criterion on a variable), this metric is not a perfect proxy for the goodness of control. For instance, the cursor might reach the target by orbiting around it and progressively getting closer (Fig. 2E, top). Although the distance continuously decreases in this case, control would not be optimal. Similarly, if the cursor is in a corner opposite to the target, any displacement will make the distance to the target decrease, irrespective of whether the trajectory is optimal (Fig. 2E, bottom). To validate that a classifier trained on cursor-to-target distance can generalize to other performance metrics, we also measured the angular error (i.e. the angle between the cursor’s velocity vector and the ideal angle to the target) and confirmed that it was higher during periods that were labeled as errors (Fig. 2F; C2: *p <* 0.001; P2: *p <* 0.001; P3: *p <* 0.001; P4: *p <* 0.001, one-sided t-test). This provides a secondary confirmation that this method of identifying errors is capable of distinguishing periods of poor and good performance.

An important feature of the error signal is that it should be detectable irrespective of the position of the target. To verify this, we replicated our analysis by performing cross-validation not across time, but across targets: for each of *n*-folds (*n* = 8), the classifier was trained solely on the trials for which the target was at 7 of the 8 possible positions, and then tested on the remaining 1 target position. This cross-validation across targets yielded qualitatively similar results (Supp. Fig. 2A-B).

However, we observed that, in most experimental sets, errors did often occur in the same direction of movement (e.g. towards the top left for the second example displayed in Fig. 2A), indicating a bias in the decoded control signal. The direction of bias differed from day to day (compare for instance Fig. 2G and Fig. 3D, which were obtained for different recording days for participant P4). Fig. 2G shows a histogram of the cursor directions during periods of correct and erroneous control for one example session, while Supp. Fig. 2C shows that the entropy of the distribution of cursor directions for all sessions is significantly lower than that of a uniform distribution, indicative of biases. One possibility is that the error detection classifier is using biased movements and decoding a directional signal instead of an actual error signal. To verify that the error classifier is not simply working as a bias direction classifier, we verified (Supp. Fig. 2D) that there are correctly detected epochs of erroneous movement and correctly ignored epochs of correct movement in every direction bin, and especially in the direction bin corresponding to the error bias (i.e. around 225° in Fig. 2G), while, if the classifier was only working based on direction, we would expect to see mostly true positives and false positives for this bin. More generally, the distribution of the cursor velocity direction in epochs flagged as erroneous by the classifier largely overlaps with that of actual erroneous control, and is not restricted to a single directional bin (e.g. compare the top right and bottom right histograms in Fig. 2G). Finally, to test the degree to which bias in the cursor kinematics impacted error detection performance, we compared, for each set, its classification accuracy to the entropy of the distribution of cursor direction during erroneous control (a lower entropy being indicative of a more biased distribution). We found no correlation between them (Fig. 2H, linear regression [35]: *T* = *−*1.279, *p* = 0.208), showing that a more biased distribution of cursor direction during faulty control does not facilitate error classification.

**Figure 3:**
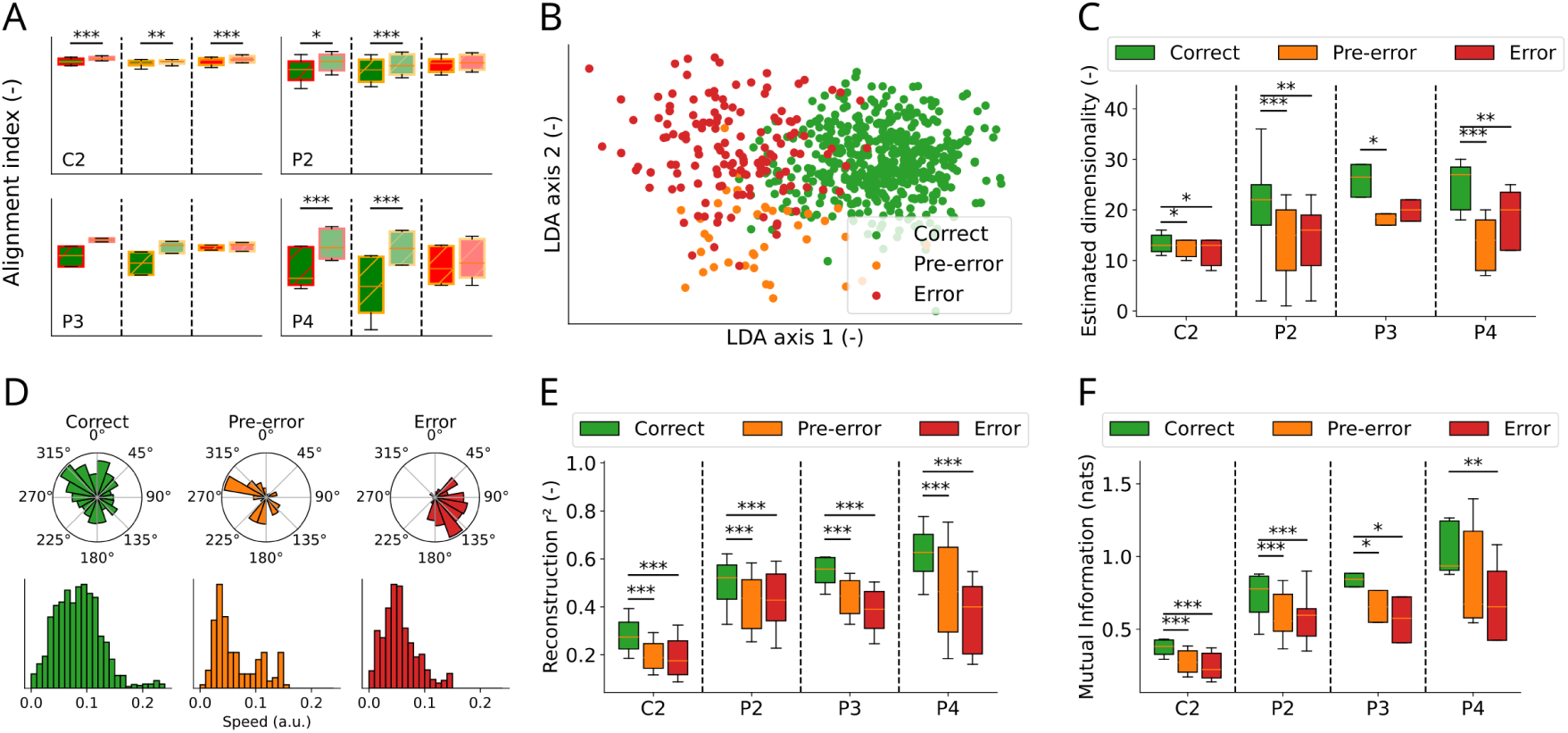
Neural activity during the pre-error window is similar to activity during erroneous control. A: Alignment index between all three subspaces (darker boxes) and their chance level obtained after shuffling (lighter boxes) for each participant (left: Correct vs. Error; middle: Correct vs. Pre-error; right: Error vs. Pre-error). Asterisks indicate alignment index values which are significantly below their chance level. B: LDA projection of neuronal activity for one example session from participant P4, showing possible separability between the *Correct*, *Pre-error*, and *Error* subspaces. C: Reduced data dimensionality is observed during epochs of erroneous and pre-erroneous control compared to the correct subspace. D: Distribution of cursor direction (top) and speed (bot-tom) during epochs of correct, pre-error and error control for one example set (consisting of several trials) from participant P4. E: Coefficient of determination between the ideal motor intent vector *v^-^_t_* (Eq. 1) and its linear reconstruction from neural activity in all subspaces. F: Mutual Information (expressed in nats) between the motor intent *v^-^_t_* and the neural activity in all subspaces.

Overall, these results indicate that a linear classifier can be trained using labeled neural features to detect epochs of erroneous movement: errors can be correctly differentiated from correct control, allowing for immediate error detection and application to real-time BCI control.

### 2.3 Neural activity during the pre-error window is similar to activity during erroneous control

Having verified that a classifier can separate periods of correct and incorrect control based on recorded neural features, we next quantify similarities and differences in the neural data between the three epochs of interest: correct control, erroneous control, and the pre-error window. We can hypothesize that the neural error signal may have two components. Firstly, there could be an exogenous component, i.e. a response to the visual feedback of the cursor moving away from the target. Secondly, there could be an endogenous component corresponding to spontaneous changes in cortical activity that lead to erroneous control. We thus propose two hypotheses regarding the nature of cortical activity in the pre-error window. The first hypothesis is that pre-error activity only corresponds to the endogenous component: since it is taking place before the actual onset of the control error, it should be free of its visual feedback. The second hypothesis is that pre-error activity is similar to error-related cortical activity and that the participant is anticipating upcoming errors. It could be possible that, although the distance to the target is still decreasing during the pre-error window, the participant already perceives that the trajectory is not optimal and likely to go off-target. A high degree of similarity between recordings in the pre-error window and erroneous control would be in favor of the second hypothesis.

A commonly used metric for assessing the similarity between two data subspaces is the Alignment Index [36, 37] (see Subspace alignment index): the index is equal to 1 if both subspaces live in the same hyperplane (i.e. if they share similar dimensions), less than 1 if they are misaligned, and 0 if they live in orthogonal planes. To assess whether two subspaces are meaningfully misaligned, we compared their respective alignment index (dark boxes in Fig. 3A) to a chance level obtained after computing the alignment index of shuffled data (lighter boxes in Fig. 3A). For 3 of 4 participants, neural activity during periods of correct control is significantly misaligned with erroneous control (i.e. has an alignment index value lower than chance level, see the green boxes in Fig. 3A. C2: *p <* 0.001; P2: *p* = 0.015; P3: *p* = 0.065; P4: *p <* 0.001, one-sided paired t-test) and pre-error neural activity (C2: *p* = 0.002; P2: *p <* 0.001; P3: *p* = 0.138; P4: *p <* 0.001, one-sided paired t-test). This is to be expected, since the classifier is precisely picking up on these differences to identify epochs of erroneous control. Performing LDA enables the visualization of the difference between these subspaces when projecting neural activity onto a lower dimensionality subspace (Fig. 3B). The same trend appears for participant P3, although not at a significant level. Additionally, we computed the alignment index between the pre-error and error subspaces (red boxes in Fig. 3A): interestingly, for 3 of 4 participants, the alignment value is not significantly different from its chance level, signifying a higher similarity between these subspaces (C2: *p <* 0.001; P2: *p* = 0.222; P3: *p* = 0.474; P4: *p* = 0.428, one-sided paired t-test). This is in line with our classification results (where we leveraged both the pre-error and error activity for error detection) and speaks in favor of the second hypothesis (that pre-error activity already includes sensory feedback, or at least a prediction of the upcoming error).

To further quantify the differences in neural activity between periods of correct, pre-error, and erroneous control, we computed the dimensionality of neural recordings during these respective epochs. Here, we define the dimensionality as the number *d* of patterns that are necessary to account for the activity of *N* recording electrodes, with *d << N* (see Dimensionality computation and [38]). The dimensionality of cortical activity has been shown to relate to behavioral data; specifically, it has been shown to be reduced during erroneous trials in cognitive tasks [39] and increased in contexts requiring fast and efficient reactions [40]. We thus expect the value of the dimensionality of motor cortical activity to be different in periods of correct and erroneous control. Indeed, we observe a lower dimensionality during erroneous epochs (Fig. 3C, *Correct* vs. *Error*; C2: *p* = 0.018; P2: *p* = 0.001; P3: *p* = 0.054; P4: *p* = 0.001, one-sided paired t-test), which is in line with the *dimensionality collapse* previously observed in the prefrontal cortex (PFC) during failed trials in cognitive tasks [39]. Interestingly, the observed dimensionality reduction tended to precede the kinematic error and was already present in the pre-error window (Fig. 3C, *Correct* vs. *Pre-error*; C2: *p* = 0.028; P2: *p* = 0.001; P3: *p* = 0.011; P4: *p <* 0.001, one-sided paired t-test), further supporting the hypothesis that pre-error activity is similar to error activity. These results are robust to the method used for assessing the dimensionality of data, since similar results were obtained when computing the Participation Ratio to estimate the number of relevant components in our data [40] (Supp. Fig. 3A). This dimensionality reduction is also in line with the smaller repertoire of movements performed during erroneous control (see the broader range of directions and speeds during correct control in Fig. 3D), which is more directionally-biased than during correct control (Supp. Fig. 2C).

To further quantify possible reductions in movement-related information during periods of erroneous control, we computed the accuracy of reconstructing the motor intent *v^-^_t_* (Eq. 1) from neural activity (see Motor intent reconstruction). *v^-^_t_* is the ideal movement direction that the cursor should follow at a given time *t* to reach the target, and can thus be used as a proxy for the BCI user’s intent [41]. The possibility to reconstruct motor intent from neural activity is reduced in both pre-error (C2: *p <* 0.001; P2: *p <* 0.001; P3: *p <* 0.001; P4: *p <* 0.001, one-sided paired t-test) and error epochs (C2: *p <* 0.001; P2: *p <* 0.001; P3: *p <* 0.001; P4: *p <* 0.001, one-sided paired t-test) as compared to periods of correct control (Fig. 3E). Furthermore, since the link between neural activity and motor intent may not be linear, we also computed their Mutual Information (MI), which is a more general measure of the dependence between two random variables (see Mutual Information); a similar trend appears, as this mutual information is significantly decreased during periods of erroneous control (Fig. 3F, green vs. red: C2: *p <* 0.001; P2: *p <* 0.001; P3: *p* = 0.013; P4: *p* = 0.004, one-sided paired t-test) as well as right before the error onset, during the pre-error window (green vs. yellow: C2: *p <* 0.001; P2: *p <* 0.001; P3: *p* = 0.011; P4: *p* = 0.256, one-sided paired t-test).

Together, these results imply that motor cortex activity occupies meaningfully non-aligned sub-spaces during periods of correct and erroneous control (Fig. 3A-B), the latter being characterized by a dimensionality reduction (Fig. 3C and Supp. Fig. 3A), and a reduction in information about motor intent (Fig. 3E-F). Crucially, these changes can already be observed before the kinematic error occurs, which further justifies leveraging neural data recorded during the pre-error window for error detection and modulation.

### 2.4 Error modulation improves participants’ performance during a 2D cursor control task

Having verified that epochs of erroneous control can be accurately detected in offline recordings, we next implemented error modulation during online BCI control in three of the participants. Error modulation was applied in parallel to motor decoding for participants C2, P2, and P4 performing the 2D cursor control task illustrated in Fig. 1A-B. When an error was detected, the decoded velocity was reduced to 30% of its computed value.

For each recording session (C2: 5 sessions; P2: 7 sessions; P4: 4 sessions), several sets of 40 trials were conducted either without (”Modulation OFF” or baseline; C2: 22 sets; P2: 11 sets; P4: 12 sets) or with (”Modulation ON”; C2: 15 sets; P2: 9 sets; P4: 11 sets) error modulation. Performance metrics were then averaged for each session and compared between both conditions.

All three participants exhibited different baseline performance (compare for instance the trajectories between C2, P2, and P4, Fig. 1B). The discrepancy between participants P2 and P4 can be attributed to differences in time since implant (P2 having been implanted nearly 10 years prior to this study) and decreasing signal quality over time for intracortical arrays [42], ultimately leading to poorer control. Despite a more recent implantation date, participant C2 had poor signal quality on one of two motor arrays (Supp. Fig. 1) that contributed little to motor decoding, likely explaining their lower performance during BCI motor tasks.

In many sessions, we observed that error modulation yielded trajectories that were straighter and more accurate (Fig. 4A and Video 1). To quantify this improvement, we computed the following metrics related to control performance: the proportion of correct trials per set (Fig. 4B), the acquisition rate (Fig. 4C), the normalized path length (Fig. 4D), the angular error (Fig. 4E), the path deviation (Fig. 4F), and the subjective difficulty reported by the participant at the end of each set (Fig. 4G).

**Figure 4:**
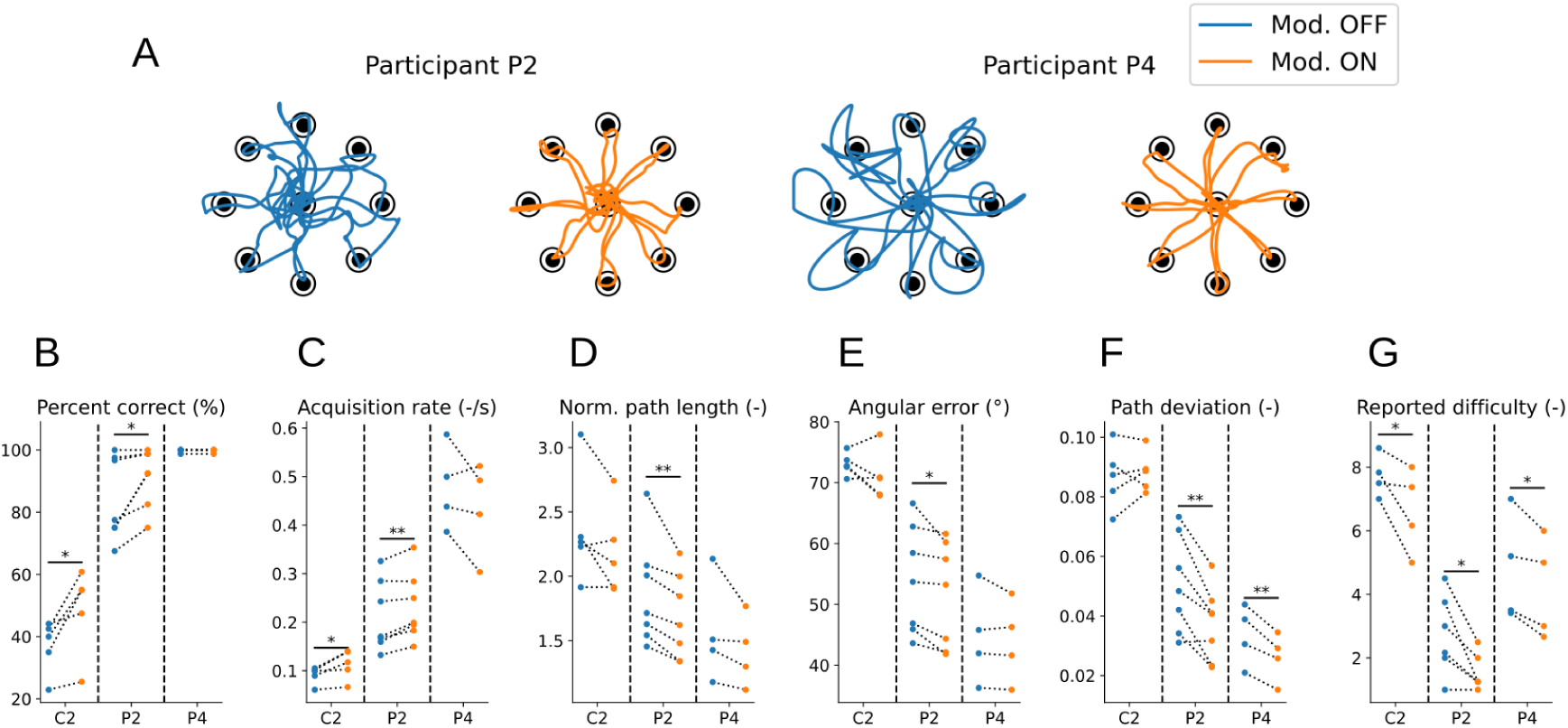
Error modulation improves participants’ performance during a 2D cursor control task. A: Trajectory examples for a block of 8 trials of the center-out and back task for participant P2 (left) and P4 (right), without (blue) and with (orange) error modulation. B-G: Performance metrics without (blue) and with (orange) error modulation. Each pair of dots represents the average performance for one session. For *Percent correct* and *Acquisition rate*, an increased value indicates that error modulation improves performances. For all other metrics, a reduced value indicates that error modulation improves performances (*Percent correct*: C2: *p* = 0.017; P2: *p* = 0.021; P4: *p* = 1. *Acquisition rate*: C2: *p* = 0.021; P2: *p* = 0.004; P4: *p* = 0.89. *Norm. path length*: C2: *p* = 0.065; P2: *p* = 0.005; P4: *p* = 0.082. *Angular error*: C2: *p* = 0.11; P2: *p <* 0.001; P4: *p* = 0.109. *Path deviation*: C2: *p* = 0.702; P2: *p* = 0.002; P4: *p* = 0.005. *Reported difficulty*: C2: *p* = 0.044; P2: *p* = 0.012; P4: *p* = 0.019, paired one-sided t-tests).

For participant C2, error modulation increased the proportion of correct trials per session (Fig. 4B), the target acquisition rate (Fig. 4C), and reduced the perceived task difficulty (Fig. 4G). However, some performance metrics (path efficiency, deviation, and angular error, Fig. 4D-F) were not improved. Importantly, error modulation improved all performance metrics for participant P2. Note that participant P4 already had very good baseline control and a nearly 100% success rate without error modulation. As such, for some metrics, error modulation does not yield a significant improvement of performance for P4 (i.e. acquisition rate, path efficiency and angular error): this is likely due to the fact that the 2D cursor control task considered here did not require fine control, hence error modulation not being required to further improve the already good baseline performance of P4.

The bottom row of Supp. Fig. 4A shows the average error probability computed by the classifier as a function of the amount of time elapsed in the trial. For participant P4, errors were mostly detected at 2 specific epochs: first, right after the onset of the trial (the cursor has a tendency to very briefly move in a random direction at the beginning of the trial before the participant corrects the trajectory, as seen for instance in the second example of Fig. 2A; this is correctly accounted for by error modulation); secondly, in the second half of the trial (when the cursor was getting close to the target and finer correction or speed reductions were performed). For participants C2 and P2, the time at which errors were detected was nearly uniformly distributed across the trial; due to their poorer control compared to P4, trajectories are less stereotypical and error modulation is more evenly distributed across the time of the trials.

The decoded speed of the cursor, i.e., the gain of the controller, has been shown to be a critical parameter for performance in BCI cursor control [13, 43]. Error modulation, in which the decoded speed is modulated depending on the output of the classifier, can be seen as a way to improve control by (dynamically) optimizing the controller gain. This is in line with previous studies, in which performance was improved either by offline gain optimization [43] or by dynamically modifying the gain of a Kalman filter [13]. One could argue that the improvements we observe here are solely due to error modulation reducing the mean speed of the cursor, hence improving the ability to accurately control it along a trajectory without actually picking up on the error signal; we believe this is unlikely, since the reduction of the average speed of the cursor induced by error modulation is often non-significant (Supp. Fig. 4B). Moreover, the classifier is correctly able to detect periods of increasing distances and higher angular errors (Fig. 2F) and to increase the target acquisition rate (Fig. 4C); it thus only reduces the cursor’s speed when it needs to be reduced.

### 2.5 Neurally-driven error modulation generalizes across tasks and contexts

Overall, error modulation yields trajectories that are more accurate, straighter, and less erratic, at the cost of slightly slower cursor control (Supp. Fig. 4B, see also the absence of improvement in target acquisition rate for participant P4 in Fig. 4C). This is to be expected, as our error modulation scheme operates by reducing the speed of the decoded velocity. This trade-off should make error modulation especially useful for tasks requiring a high level of precision and stability rather than fast displacements (such as the 2D target reaching task used previously). To verify this idea and to assess whether an error classifier can robustly generalize across tasks and contexts, we trained an error classifier using the same center-out task as in the previous sections, and then tested them on two different tasks: a “click-and-drag” cursor control task (Fig. 5A), and a suite of tasks from the 2024 Cybathlon BCI competition (Fig. 5C).

**Figure 5:**
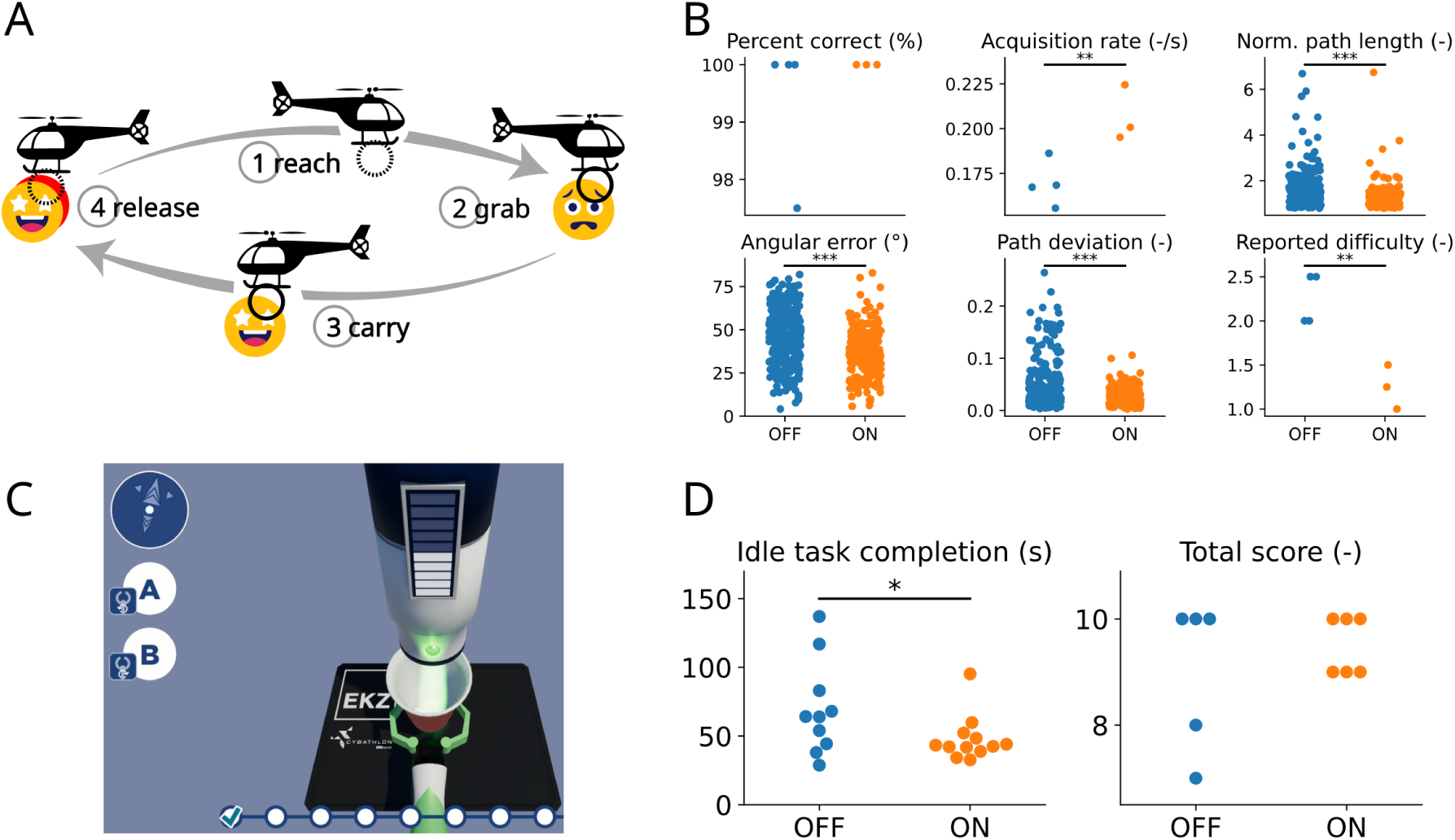
Neurally-driven error modulation generalizes across tasks and contexts. A: Schematics of the grasp and drag task, gamified in the form of a “helicopter rescue” task and performed by participant P2. The participant moved the cursor to the target (*reach* phase), clicked to grab it while remaining close to it (*click* phase), then dragged it back to the center (*center* phase) before releasing it (*release* phase). Redrawn from [6]. B: Performance metric improvement for participant P2 due to error modulation (*Percent correct*: *p* = 0.218; *Acquisition rate*: *p* = 0.008; *Norm. path length*: *p <* 0.001; *Angular error*: *p <* 0.001; *Path deviation*: *p <* 0.001; *Reported difficulty*: *p* = 0.002, one-sided t-tests). C: Illustration of the idle task from the Cybathlon BCI race [33] performed by participant P4. The participant needs to control the arm holding the cup and maintain it idle under the ice dispenser until it is filled to complete the task. Image: CYBATHLON 2024 BCI Race Game by CYBATHLON, ETH Zurich and Koboldgames GmbH, Switzerland. D: Completion time for the idle task (left; *p* = 0.032, one-sided t-test) and total score for the full BCI race (right; *p* = 0.221, one-sided t-test), without (blue) and with (orange) error modulation, for participant P4.

First, we applied error modulation to a “click-and-drag” variation of the 2D cursor control task [6], in which the participant not only has to move a cursor to a target, but also needs to grab the target and release it at the center of the screen, akin to a click-and-drag action performed with a mouse (Fig. 5A). This task requires the participant to maintain the cursor position over the target area to click the target, and similarly to remain stable in the center area to release it. Click and release actions were decoded in parallel to cursor velocity using grasp-related neural transients [6]. The task was performed by participant P2. For this more demanding version of the cursor control task, error modulation also yielded improved performance with a significantly higher target acquisition rate and a lower reported difficulty (Fig. 5B), demonstrating its usefulness for tasks in which accuracy is critical.

Finally, to further validate the robustness of error modulation to task and context, we applied it to the BCI tasks from the recent Cybathlon competition [33]. The Cybathlon BCI race includes different realistic tasks performed in a video game environment: controlling a wheelchair in a room, a robotic arm to operate a machine or open a lock, and a cursor to click on a specific area of the screen. These tasks are completed by controlling the velocity of a video game actuator (either a cursor or an avatar of a prosthetic arm depending on the task) using the same decoding scheme as in the previous sections. Of particular interest is the idle task, where the participant needs to control a robotic arm to bring a cup under an ice dispenser and maintain it stable until the cup is filled (Fig. 5C). This task was performed by participant P4. We found that error modulation reduced completion times during the idle task (Fig. 5D, left). To ensure error modulation did not compromise performances in the other tasks of the BCI competition, we also measured each race’s score (i.e. the number of successfully completed tasks out of ten), which provides a better estimate of the control performance during a race than its completion time since failing a task will decrease the race completion time. We verified that error modulation did not lead to reduced race scores (Fig. 5D, right). Note that each race contains two idle tasks, hence the two-fold difference in the number of data points in both panels of Fig. 5D.

For both tasks (i.e. the helicopter rescue task, and the Cybathlon BCI tasks), the error classifier was trained on 2D center-out data, as in the previous sections, and not on task-specific data; this further hints at the generalizability of the classifier across tasks and contexts.

## 3 Discussion

Brain-computer interfaces rely on accurate decoding of motor intent from neural population activity. However, control is rarely perfect, falling short of able-bodied performance [5, 7]. The ability to estimate intent is limited because BCIs rely on noisy recordings from a limited number of electrodes [10] and the neural population activity can be impacted by changes in task context, mental state, or recording instabilities [15, 42, 44–46]. This calls for novel approaches for improving BCI accuracy and usability. Using a neural signature of ongoing errors, we trained a classifier to detect periods of erroneous control from cortical activity; this enabled online error detection during BCI control to minimize errors without any specific action from the participant. Our contributions are three-fold. First, we show that neural signatures of ongoing errors can be detected earlier than previously thought [18, 19, 25, 29, 32]. Analysis suggests that participants may be predicting that their trajectory is going to go off course before the error actually manifests. Second, we show that this can be leveraged to perform early error detection and prevention with human BCI users, yielding significant improvements in various performance metrics including success rate and perceived difficulty. Unsurprisingly, the effect of error modulation on performance was greatest when the baseline level of performance was poorer. Finally, we show that error modulation can generalize to more complex tasks and contexts without task-specific training.

Our results reveal neural correlates of errors in motor cortex during iBCI tasks. This is in line with previous studies using other recording modalities or tasks, or in NHPs. More specifically, previous studies have already suggested that the neural error signal could be used to improve the accuracy of BCIs [20, 22, 25, 26, 29, 30, 32, 47], often in the context of motor BCIs, but also for different applications such as text spelling [21] or speech production [32]. In a recent study [31], an error signal was detected using intracranial electroencephalography (iEEG) from the insula of human participants, and used to perform error correction in a binary classification task. However, the error classifier was trained on long windows of data (500ms) studied 1s after the onset of error, making this modality likely unsuitable for error correction in continuous control tasks.

Interestingly, we were able to detect an error signal in neural activity immediately preceding the kinematic error, suggesting that participants may be predicting trajectory deviations before they occur. In our case, and in previous work, we defined an error as an increasing distance between the cursor and the target. Our analysis (section 2.3) suggests a similarity between neural activity during actual erroneous control and during the pre-error window, suggesting that this prediction can be detected earlier than previously thought.

We demonstrated that error modulation can improve BCI control on a number of metrics in participants with varying degrees of baseline control. Importantly, the subjective difficulty was improved for all participants, who reported, for instance, that “This decoder [with error modulation] was my favorite. It felt easier” (participant P2) or “I felt I had better control of the cursor” (participant P4). For participant C2, significantly improved performance metrics also include the task success and target acquisition rates, but not the other metrics. This is in line with previous studies involving NHPs, in which the improvement was not significant for all metrics and all subjects (namely, it was only significant for the orbiting time, but not for the total time to target [18]). We suspect that C2 may have been limited by a lower overall signal quality and yield as compared to the other participants. We have hence shown that an error signal can be detected both in offline (Fig. 2) and online data (Fig. 4), and could still be decoded a long time after implantation (nearly 10 years for participant P2), even with limited signal quality.

We presented some evidence that this approach will generalize to more complex tasks. Previous studies had shown that, during a BCI task (either controlling a cursor on a screen or a prosthetic limb), remaining idle can be challenging and non-adaptive decoders often exhibit poor performances at low velocities [13, 48, 49]. Our results show that error modulation is especially efficient when stability and accuracy are required (section 2.5). Furthermore, the error classifier used for error modulation in the helicopter rescue task (Fig. 5A) and the Cybathlon tasks (Fig. 5C) was trained using data from a simple 2D cursor control task (Fig. 1). These results show that the error signal detected during one task can generalize to another where the visual feedback of the error may be different than the task in which it was trained.

Our study has some limitations. Namely, the classifier pipeline does not allow for perfect epoch classification on offline data (Fig. 2), nor does it allow users to achieve perfect performance during online BCI use (Fig. 4). Furthermore, the classifier requires recalibration at the beginning of each day, just like the movement decoder algorithm. An interesting future step would be to not only classify periods of correct and erroneous control, but to also actually predict the direction and amplitude of the error from neural activity. This would allow for not simply damping, but rather correcting errors on-the-fly in the desired direction, at the risk of a higher performance cost in case of faulty prediction (which could further aggravate the directional error). A recent study [27] showed that positional error can be decoded from offline EEG recordings using a convolutional classifier. However, the possibility of performing active error corrections during online BCI control still needs to be assessed. Interestingly, those results were obtained with a modification to standardized EEG electrodes positions: two of them were relocated close to the cuneus and precuneus, which are known to encode visual information [50, 51]. In comparison, intracortical electrode arrays which are used for implanted BCIs (including the present study) are usually implanted in the motor or premotor areas [52] which may convey less information about visual feedback (although ECoG data have shown the presence of error-related signals in several brain areas, including the somatosensory, motor, premotor, and parietal areas [29]).

To conclude, detecting the error signal enabled on-the-fly error detection and damping to improve BCI motor decoding performance. We identified early neural signatures of control errors and improved control by modulating the decoded output on multiple computer-based BCI tasks. This addresses a significant priority for BCI users by paving the way towards more reliable control.

## 4 Material and methods

### 4.1 Participants

Four male participants (C2, P2, P3, and P4) in an ongoing BCI clinical trial participated in the experiments described here. C2 was in his 60s at the time of implant with a C4 ASIA D spinal cord injury and a brachial plexus injury 4 years prior. P2 was in his 20s at the time of implant with a C5 ASIA B spinal cord injury 10 years prior. P3 was in his 20s at the time of implant with a C6 ASIA A spinal cord injury 12 years prior. P4 was in his 30s at the time of implant with a C4 ASIA A spinal cord injury 11 years prior. C2 has very limited movement of his upper limb and hand. P2 and P3 have residual arm and wrist movement, but no hand function. P4 has no volitional movement below the neck. All four participants were implanted with intracortical microelectrode arrays (”Neuroport Arrays”, Blackrock Neurotech, Salt Lake City, UT, USA). Two arrays were implanted in the precentral gyrus of the motor cortex, while the other two arrays were implanted in the somatosensory cortex. Data from the somatosensory cortex were not analyzed in this study. P2 had 88 electrodes in each motor cortex array, while C2, P3 and P4 had 96 channels in each motor cortex array.

This study was conducted under an Investigational Device Exemption (IDE) from the Food and Drug Administration and approved by the Institutional Review Board of the University of Pittsburgh. This study was completed as part of a clinical trial registered at clinicaltrials.org (NCT01894802). Informed consent was obtained from all participants prior to the beginning of their involvement in the study and all study procedures comply with the Declaration of Helsinki.

### 4.2 Neural Recordings

Recordings were obtained with the Blackrock Neurotech Neuroport System and preprocessed following the same methodology as in a previous study [6]. Briefly, neural signals were recorded on the 2 intracortical electrode arrays implanted in the precentral gyrus of the motor cortex. Signals were filtered using a 1st order 750 Hz high-pass filter [53], logged as threshold crossings (−4.5 RMS), and binned at 50 Hz. The binned counts were then convolved with a 440 ms decaying exponential filter to provide a smoothed estimate of firing rate for input to the BCI decoder.

### 4.3 Movement decoding

The dimensionality of the binned spike counts was reduced to 20 components using factor analysis and the resulting components were used to train a Kalman filter for velocity decoding. In line with previous studies [6, 27], the velocity decoder was calibrated in 2 steps. In the first step (referred to as *observation calibration*), the cursor moved automatically to the target, and participants were instructed to follow it by performing an imagined movement using their preferred imagery [52]. The recorded neural activity and the corresponding cursor velocity were used to calibrate a first version of the decoder. In the second step (referred to as *closed-loop calibration*), this decoder was used by the participant to control the cursor such that all commands orthogonal to the ideal direction were attenuated, thus limiting movement to a one-dimensional line towards the target. Data from the closed-loop calibration was used to calibrate a new decoder that was then used for 2D BCI control. Due to potential changes in the recorded neural activity across sessions [44], a new decoder was trained each day.

### 4.4 Error definition

In line with previous studies [18], we defined periods of correct and erroneous BCI control using the instantaneous distance between the current position of the cursor and the target (Fig. 1C). At each time step *t*, the *motor intent* vector *v^-^_t_* is defined as the difference between the position of the current target [*x^-^_t_, y^-^_t_*] and the current cursor position [*x_t_, y_t_*]:

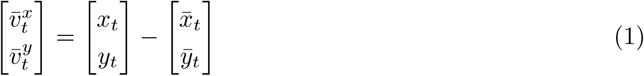

In case of ideal control, the absolute distance from the target 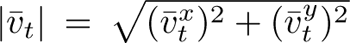 should continuously decrease. However, in practice, periods of erroneous control are defined as the periods during which this distance between cursor and target is increasing (red in Fig. 1C) instead of decreasing (correct control, green in Fig. 1C). Finally, the *pre-error* window was defined as the 200ms window immediately preceding the onset of a spontaneous error (yellow in Fig. 1C).

### 4.5 Error classification

Our approach to train the error classifier was as follows. First, we examined firing rates and kinematics using 80 ms non-overlapping sliding windows (i.e. 4 samples at 50 Hz). Then, each sample was labeled based on the change in position of the cursor relative to the target (Fig. 1C): if the cursor-to-target distance was decreasing while the features were recorded, the sample was labeled as *correct control* (green); if it was increasing, it was labeled as *erroneous control* (red). Finally, these labeled samples obtained during a first online control step are used to train a naive Bayes classifier, which is then used during subsequent online control steps. Training both the movement decoder (see Movement decoding) and the error classifier thus requires 3 consecutive steps: 40 trials of *observation calibration*, 40 trials of *closed-loop calibration* (which are used to train the final movement decoder), and 40 trials of online control (from which the data used to train the error classifier are obtained). Whenever the probability of erroneous control computed by the classifier reaches a certain threshold *τ*, error modulation was activated by reducing the cursor’s speed to 30% of its computed value.

An important element of the classifier’s design is the limitation of false alarms (i.e. implementing error modulation while the trajectory is actually correct). These would be particularly frustrating for the user and could decrease performance [31]. The false alarm rate is controlled by the threshold parameter *τ*: if the error probability computed by the classifier is above *τ*, the sample is classified as erroneous. To optimally set this hyperparameter, we computed the classification accuracy (as in Fig. 2B) averaged across sessions as a function of *τ* (Supp. Fig. 5A). Interestingly, this accuracy appears to be mostly independent of *τ*: indeed, the distribution of probability values computed by the classifier appears to be bimodal, mostly taking values close to 0 or 1 (Supp. Fig. 5B for an example session). Any value of *τ* in [0, 1] would yield similar results. For consistency, the same value *τ* = 0.85 was used across all trials and participants.

### 4.6 Performance metrics

For each recording session (C2: 5 sessions; P2: 7 sessions; P4: 4 sessions), several sets of 40 trials were conducted either without (”Modulation OFF”) or with (”Modulation ON”) error modulation. The first set of each session was always without error modulation (since a first set of online control is required to obtain the data for training the error classifier, see Error classification). Sets with and without error modulation were then alternated until the end of the recording session. Participants were not explicitly told that different decoders were used, nor that error modulation was tested; however, due to error modulation modifying the aspect of the cursor’s trajectory, participants could report differences between both conditions (i.e. whether modulation was on or off).

Recording sessions had variable lengths (some having only 1 set for each condition): alternating conditions instead of randomizing them ensured a comparable number of sets was recorded for each condition. For some sessions, additional sets without error modulation were performed and included at the end of the session, yielding a higher number of sets in the “Modulation OFF” condition (C2: 22 sets; P2: 11 sets; P4: 12 sets) than in the “Modulation ON” condition (C2: 15 sets; P2: 9 sets; P4: 11 sets).

Different metrics were used to assess the participants’ performance during online BCI control:

The **success rate** (-) corresponds to the number of successful trials (out of 40) for each set. A successful trial is defined as the completion of a successful *reach* phase and a successful *center* phase.

The **acquisition rate** (*s^−^*^1^) corresponds to the number of successful trials in a set normalized by its total duration.

The **normalized path length** (-) corresponds, for each trial phase (i.e. *reach* and *center*, both being included in the analysis), to the length of the trajectory followed by the cursor normalized by the length of the ideal trajectory (a straight line between the center and the target). It was computed only for successful phases (i.e. where the target was reached).

The **angular error** (*^◦^*) was computed at each time step as the angle between the decoded velocity vector and the ideal velocity vector (that links the current cursor position and the target). This value was averaged over all time steps to obtain a single value per trial phase.

The **path deviation** (-) is the average value, for each trial phase, of the normal distance of the cursor from the ideal trajectory. It was computed only for successful phases.

At the end of each set of 40 trials, participants were asked to rank the perceived **reported difficulty** (-) on a scale from 0 (very easy) to 10 (very hard).

In Fig. 4 (target reaching task), metric values were averaged over each recording sessions. In Fig. 5B (helicopter rescue task), fewer sessions were obtained due to the task being longer; metric values were thus averaged per sets (success rate, acquisition rate, reported difficulty) or trials (normalized path length, angular error, path deviation).

### 4.7 Dimensionality computation

The embedding dimensionality of a dataset [38] corresponds to the volume occupied by the dataset, or to the number of components or dimensions that are required to describe it. Although neural activity is recorded from a set of *N* electrodes, covariance patterns between electrodes’ activity actually constrain neural activity to a hyperplane of dimensionality *d << N*, *d* being its embedding dimensionality.

In line with previous studies, we used two metrics to approximate this dimensionality: first, we used the number of principal component analysis (PCA) components that are required to explain 80% of the data’s variance [54]; this estimated value of the dimensionality is reported in Fig. 3C.

Second, to ensure results were robust to the method used for estimating the dimensionality, we computed the Participation Ratio (PR), which is a measure of the spread of the eigenvalues of the data’s covariance matrix [40]:

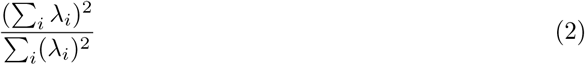

where *{λ_i_}1≤i≤N* are the eigenvalues of the data’s covariance matrix. This value is reported in Supp. Fig. 3A.

### 4.8 Subspace alignment index

The alignment index *H_A,B_* between two datasets *A* and *B* is computed as such: first, an orthonormal basis of the first dataset *A* is obtained by performing PCA with *M* components. Then, the alignment index is computed as the fraction of the variance of the second dataset *B* that is explained by this basis:

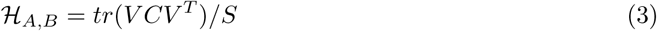

where *V* is the *M*-by-*N* matrix corresponding to the orthonormal basis of the first dataset *A*, *C* is the *N*-by-*N* covariance matrix of the second dataset *B*, and *S* is the sum of the first *M* singular values of *C*. Since the definition of the alignment index is not symmetrical (i.e. *H_A,B_ ≠ H_B,A_*), the index *H* that we show in Fig. 3A is a symmetrical version of Eq. 3:

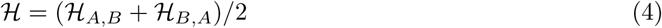

Results in Fig. 3A were obtained for *M* = 10, although different values for *M* yield similar results (Supp. Fig. 3B).

The index is thus equal to 1 if the datasets live in the same hyperplane, and to 0 if they live in orthogonal hyperplanes. But even datasets sampled from the same subspace would not be perfectly similar and could hence have an alignment index lower than 1 because of noise or undersampling, for instance. To assess whether two subspaces *A* and *B* are meaningfully non-aligned, we computed a chance alignment level which measures how sub-datasets from the same condition are expected to be aligned. To avoid sample-to-sample correlation, for each recording session, the dataset formed by joining *A* and *B* was split into 40 chunks of equal size, half of them forming a sub-dataset *Ã* and the other half forming a sub-dataset *B^∼^*. The alignment index between *Ã* and *B^∼^* is the control chance level. An alignment index (Eq. 4) lower than chance level indicates that both datasets are more misaligned than chance.

### 4.9 Mutual Information

Mutual Information (MI) is a nonlinear measure of the correlation between two variables. The mutual information *I*(*X*; *Y*) between two random variables *X* and *Y* (expressed in nats) quantifies the amount of information about the latter that is carried by the former, and can be computed as the sum of their entropies *H*(*X*) and *H*(*Y*) minus the entropy of their joint distribution:

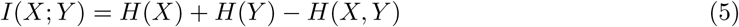

It is equal to 0 in the case of independence, and increases as *X* and *Y* are related. In our case, *Y* is an *N*-dimensional variable corresponding to motor cortex activity, while *X* is 2-dimensional and corresponds to the motor intent (Eq. 1). If *X* follows a multivariate Gaussian distribution, its entropy can be analytically computed as

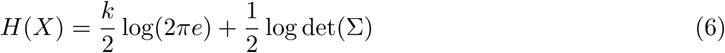

where *k* is the number of components in *X* and Σ is its covariance. To compute the mutual information between the neural activity and the motor intent, we made the approximation that these variables (as well as their joint distribution *X, Y*) were normally distributed and computed their covariance matrices from observations. Then, the entropies of *X*, *Y*, and of the joint *X, Y* were computed using Eq. 6. Finally, these values were used to approximate the mutual information (Eq. 5).

### 4.10 Motor intent reconstruction

At each time step *t*, the *motor intent* vector *v^-^_t_* was defined as the difference between the position of the current target [*x^-^_t_, y^-^_t_*] and the current cursor position [*x_t_, y_t_*] (Eq. 1). This difference between the target position and the cursor position (called the *Position Error* in [41] and interpreted as a linear approximation of the user’s feedback control policy) corresponds to the ideal trajectory towards the target and can heuristically be used as a proxy for the user’s intent. Although different methods for defining user intent exist [55], they ultimately yield the same BCI decoding accuracy [41].

For a recording session of *T* data points, these values can be concatenated into a *T*-by-2 matrix *X*. Similarly, neural recordings can be concatenated into a *T*-by-*N* matrix *Y* (*N* being the number of recording electrodes). The former can be linearly reconstructed from the latter:

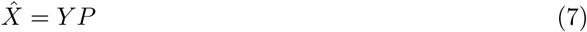

with

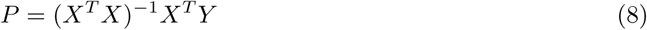

The difference between the actual motor intent *X* and the linearly reconstructed one *X^^^* is quantified by computing the Pearson correlation coefficients between their respective *x* and *y* dimensions and averaging both values.

## Acknowledgments

Research reported in this publication was supported by the Swiss National Science Foundation under grant number P500PM 210800, the University of Pittsburgh Hunter Family Foundation Innovation in Neuroscience Program, and by the National Institute Of Neurological Disorders And Stroke of the National Institutes of Health under Award Numbers R01NS121079 and UH3NS107714. The content is solely the responsibility of the authors and does not necessarily represent the official views of the National Institutes of Health. We thank Robert A. Gaunt and Lee E. Fisher for the fruitful discussions, and Isabelle Com for the illustration.

## 5 Supplementary material

**Video 1**: Trajectory during example trials from participant P2, without (blue, left) and with (orange, right) error modulation. Trajectories are accelerated 10 times.

---

**Supplementary Figure 1:**
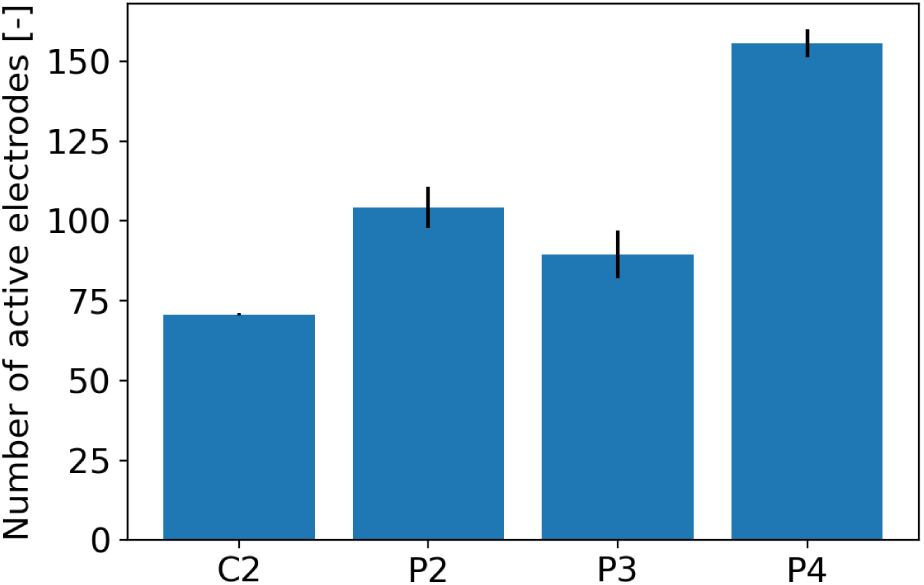
Number of included electrodes (defined as the number of electrodes in both arrays for which the firing rate is above the set mean at least 5% of the set duration), averaged over all sets for each participant.

**Supplementary Figure 2:**
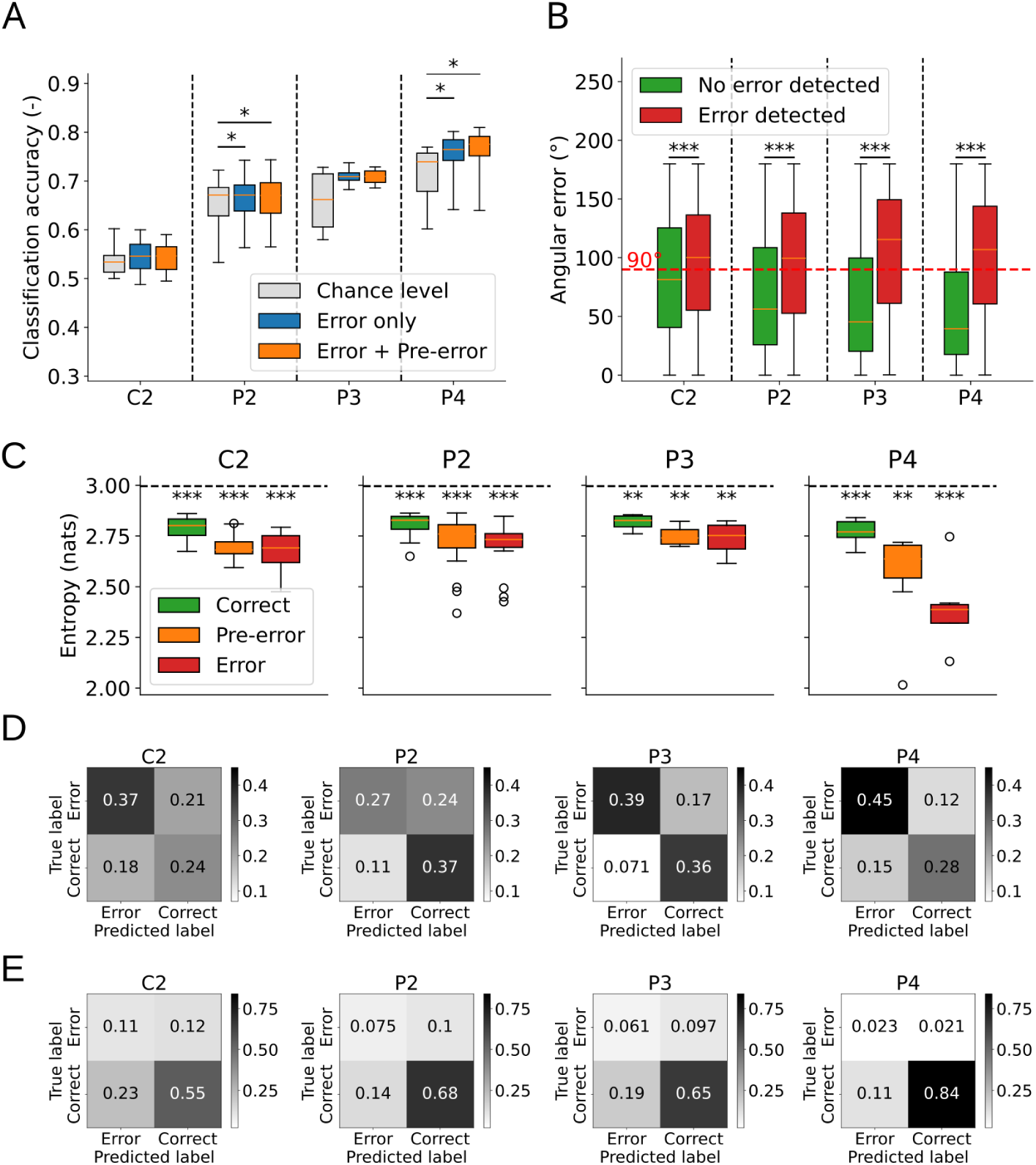
A: Same results as in Fig. 2B but with cross-validation performed across targets (*Chance level* vs. *Error only*: C2: *p* = 0.16; P2: *p* = 0.015; P3: *p* = 0.092; P4: *p* = 0.038, one-sided paired t-test. *Chance level* vs. *Error + Pre-error*: C2: *p* = 0.194; P2: *p* = 0.017; P3: *p* = 0.119; P4: *p* = 0.018, one-sided paired t-test). B: Same results as in Fig. 2F but with cross-validation performed across targets (C2: *p <* 0.001; P2: *p <* 0.001; P3: *p <* 0.001; P4: *p <* 0.001, one-sided t-test). C: Entropies of the distribution of the cursor velocity direction in the correct, pre-error, and error windows, for all sets from each participant. The black dashed line represents the entropy of a uniform distribution, which is significantly higher than observed entropies (C2: *p <* 0.001, *p <* 0.001, *p <* 0.001; P2: *p <* 0.001, *p <* 0.001, *p <* 0.001; P3: *p* = 0.002, *p* = 0.002, *p* = 0.006; P4: *p <* 0.001, *p* = 0.002, *p <* 0.001, one-sample one-sided t-tests). D: Confusion matrices (averaged over all sets for each participant) for the direction bin containing the most erroneous epochs in each set (e.g. 225*^◦^ −* 270*^◦^* for the set illustrated in Fig. 2G). E: Same results for the direction bin containing the least erroneous epochs in each set (e.g. 0*^◦^ −* 45*^◦^* for the set illustrated in Fig. 2G).

**Supplementary Figure 3:**
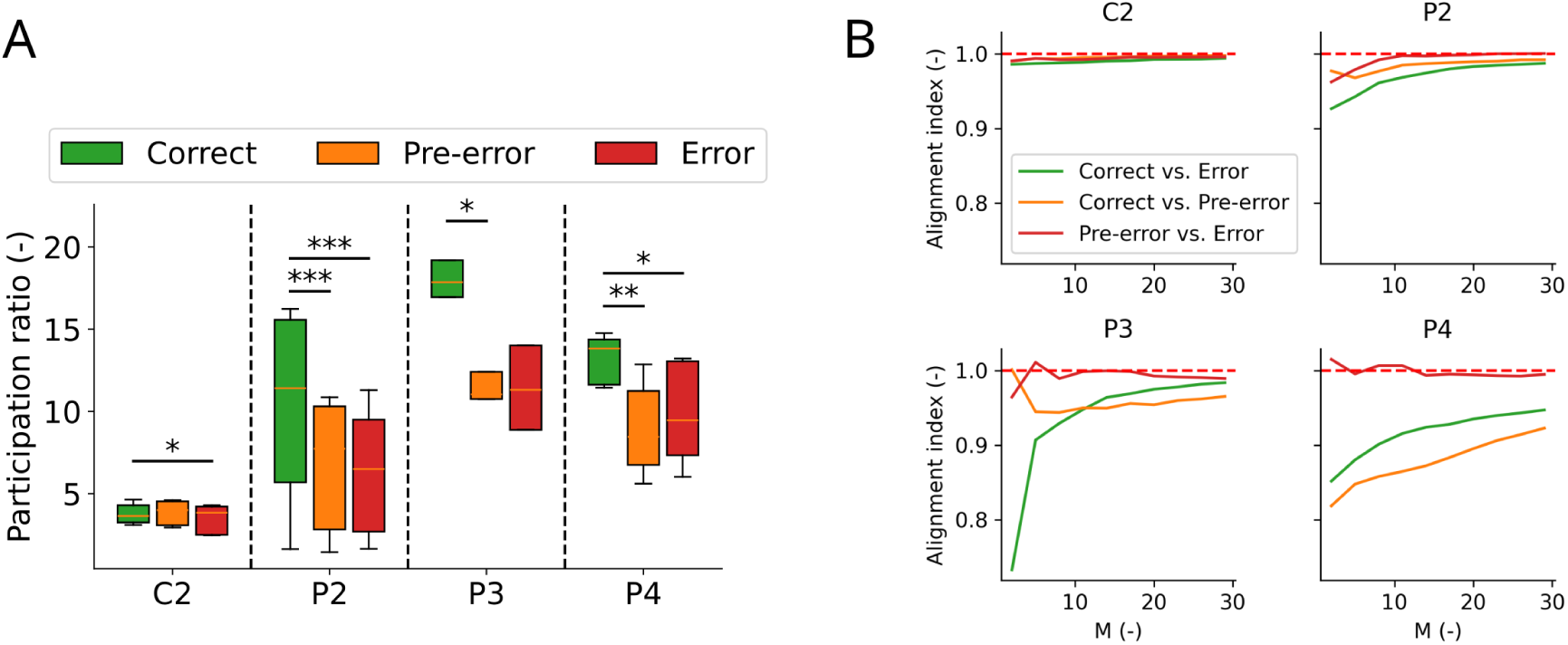
A: Same results as in Fig. 3C but using the participation ratio (see Dimensionality computation) for estimating the dimensionality of each subspace. *Correct* vs. *Error*: C2: *p* = 0.033; P2: *p <* 0.001; P3: *p* = 0.074; P4: *p* = 0.031, one-sided paired t-test. *Correct* vs. *Pre-error*: C2: *p* = 0.414; P2: *p <* 0.001; P3: *p* = 0.042; P4: *p* = 0.005, one-sided paired t-test. B: Same results as in Fig. 3A but for different values of the parameter *M* (see Subspace alignment index).

**Supplementary Figure 4:**
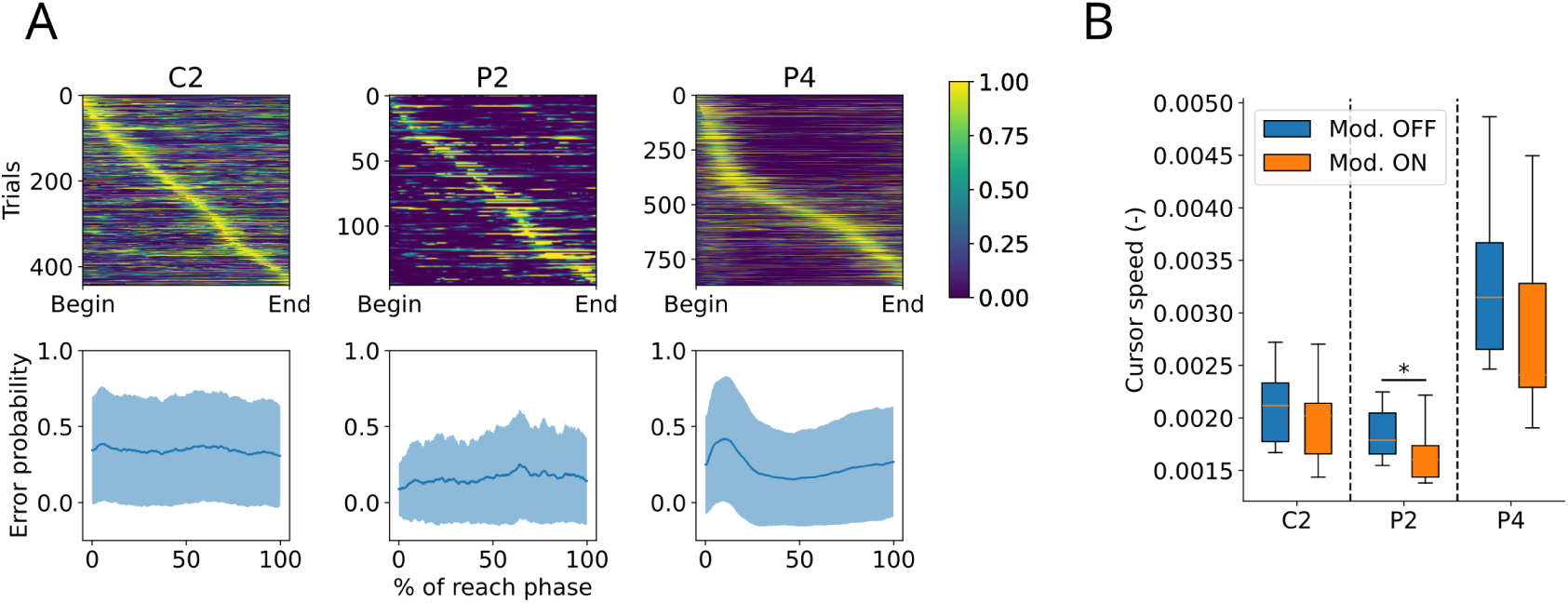
A: Top row: Heatmap showing the error probability computed by the classifier for different trials (vertical axis) as a function of the trial completion (horizontal axis). Trials are ordered by peak error probability timepoint. Bottom row: same results, averaged over all trials. Shaded area: standard error. B: Cursor speed for all phases of the trials studied in section 2.4 for all three participants (C2: *p* = 0.116; P2: *p* = 0.042; P4: *p* = 0.1, one-sided t-test).

**Supplementary Figure 5:**
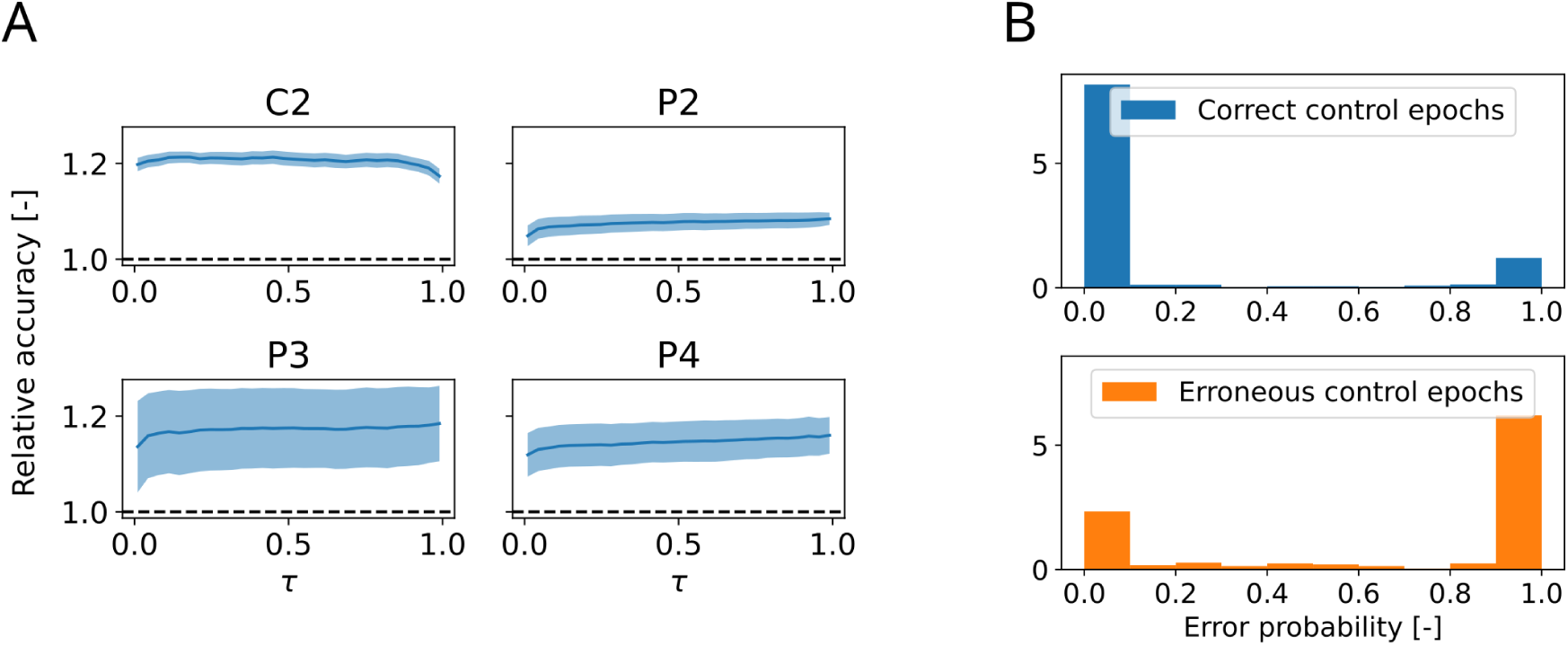
A: Average classification accuracy (normalized by the chance level as defined in section 2.2) for all sessions of each participant as a function of the error detection threshold *τ*. Shaded area: standard error of the mean. B: Histograms of the error probability values computed by the classifier during epochs of correct (top) and erroneous control (bottom) for an example set from participant P4.

**Supplementary Table 1:** Experimental sets of 40 trials each used in sections 2.2 and 2.3. Durations and proportions of erroneous control are averaged over all sets for each participant.

| Participant | Number of sets | Duration | Proportion of erroneous control |
| --- | --- | --- | --- |
| C2 | 16 | $245.68 \pm 34.49\text{s}$ | $50.11 \pm 3.74 \%$ |
| P2 | 17 | $260.23 \pm 86.42\text{s}$ | $37.93 \pm 4.76 \%$ |
| P3 | 4 | $197.0 \pm 8.53\text{s}$ | $36.48 \pm 6.43 \%$ |
| P4 | 7 | $158.95 \pm 87.9\text{s}$ | $30.46 \pm 5.87 \%$ |

## Notes

https://github.com/pitt-rnel/error_signal_analysis

### Competing Interest Statement

JW and BD served as consultant for Blackrock Neurotech, Inc., the company that sells the microelectrode arrays and data acquisition system used in this study. JLC has previously received research funding from Blackrock Neurotech.

